# Postnatal mechanical loading drives adaptation of tissues primarily through modulation of the non-collagenous matrix

**DOI:** 10.1101/2020.04.23.058081

**Authors:** D. E. Zamboulis, C. T. Thorpe, Y. Ashraf Kharaz, H. L. Birch, H. R. C. Screen, P. D. Clegg

**Affiliations:** Institute of Ageing and Chronic Disease, Faculty of Health and Life Sciences, University of Liverpool, Liverpool, L7 8TX, United Kingdom; Comparative Biomedical Sciences, The Royal Veterinary College, Royal College Street, London, NW1 0TU, United Kingdom; University College London, Department of Orthopaedics and Musculoskeletal Science, Stanmore Campus, Royal National Orthopaedic Hospital, Stanmore, HA7 4LP, United Kingdom; Institute of Bioengineering, School of Engineering and Materials Science, Queen Mary University of London, London, E1 4NS, United Kingdom

## Abstract

Mature connective tissues demonstrate highly specialised properties, remarkably adapted to meet their functional requirements. Tissue adaptation to environmental cues can occur throughout life and poor adaptation commonly results in injury. However, the temporal nature and drivers of functional adaptation remain undefined. Here, we explore functional adaptation and specialisation of mechanically loaded tissues using tendon; a simple aligned biological composite, in which the collagen (fibre phase) and surrounding predominantly non-collagenous matrix (matrix phase) can be interrogated independently. Using an equine model of late development, we report the first phase-specific analysis of biomechanical, structural and compositional changes seen in functional adaptation, demonstrating adaptation occurs postnatally, following mechanical loading, and is almost exclusively localised to the non-collagenous matrix phase. These novel data redefine adaptation in connective tissue, highlighting the fundamental importance of non-collagenous matrix and suggesting that regenerative medicine strategies should change focus from the fibrous to the non-collagenous matrix phase of tissue.

## Introduction

Functional adaptation of load-bearing tissues such as tendon is crucial to ensure the tissue is specialised appropriately to meet functional needs. Adaptation to mechanical requirements is key in healthy development and homeostatic tissue maintenance, with poor tissue optimisation during maturation likely a key contributor to increased injury risk later in life. Dysregulated homeostasis and long-term under- or over-stimulation leads to maladaptation, changes in tissue integrity, and reduced mechanical competence and is implicated in the disease aetiology of load-bearing tissues(Freedman et al., 2015; Gardner et al., 2008). Understanding the developmental drivers of structural specialisation and their association with mechanobiology is thus of fundamental importance for healthy ageing and disease prevention in musculoskeletal tissues(R. Choi et al., 2018; Chavaunne T. Thorpe et al., 2013). Such knowledge will help identify future targets for therapeutic interventions, and thus address the current lack of effective musculoskeletal disease treatments with new, evidence-based approaches to disease management. However, there is currently little knowledge of the key extracellular matrix (ECM) components associated with structural specialisation, the temporal nature of their adaptation, or the stimuli that drive adaptation.

As the principal structural component of connective tissues, collagen expression at the gene and protein level has been the focus of the majority of studies in relation to loading, with some studies reporting increases in collagen synthesis and others noting collagen degradation in response to loading, depending on the tissue function or tissue structure in different species(R. Choi et al., 2018; Magnusson & Kjaer, 2019). In materials science, this collagen structural framework is commonly referred to as the “fibre phase” of a material, surrounded by the primarily non-collagenous and glycoprotein-rich components of the ECM, termed the “matrix phase”(Armiento et al., 2018; C. T. Thorpe & Screen, 2016). This distinction is important, as it describes a fibre composite material, in which “fibre” and “matrix” phases have different mechanical properties, and overall tissue mechanical properties and function are governed by the interplay of these two phases. When looking to understand functional adaption of a tissue, it is necessary to look at adaption of all ECM components.

Whilst fibre phase adaption has received some attention, adaptation of the matrix phase to mechanical stimuli remains poorly defined. Indeed, it is notable that the numerous studies investigating the mechanoresponsive nature of load-bearing tissues tend to restrict their focus to specific fibre or matrix components with no spatial distinction, and also focus on a single element of either structural or mechanical adaptation, such that limited information is gained(Cherdchutham, Becker, et al., 2001; R. K. Choi et al., 2019; Mendias et al., 2012). In order to provide the necessary complete profile of adaptive behaviour, it is crucial that phase-specific, temporospatial adaptation in the context of both structure and function is defined.

Identifying the drivers of adaptation requires use of a model system in which the temporospatial nature of adaptation can be fully profiled. Tendon provides the ideal model for such a study. It is well-established that mature tendons can present structural and mechanical specialisms(Chavaunne T. Thorpe et al., 2015; Chavaunne T. Thorpe, Karunaseelan, et al., 2016; Chavaunne T. Thorpe, Riley, et al., 2016) and be grouped into two clear functional groups; stiff positional tendons simply connect muscle to bone to effectively position limbs, whilst elastic energy-storing tendons increase locomotor efficiency by stretching and storing energy which they return to the system on recoil(Batson et al., 2010; C. T. Thorpe & Screen, 2016; C T Thorpe et al., 2012). Further, the simple aligned organisation of tendon means that fibre and matrix phases are spatially distinct, enabling structural and mechanical characterisation of each phase independently, by comparing fascicles (fibre phase) and inter-fascicular matrix (IFM; matrix phase)(C. T. Thorpe & Screen, 2016; C T Thorpe et al., 2012). Finally, use of equine tendon provides access to an exceptional model of adaptation. The equine superficial digital flexor tendon (SDFT) has been shown to be highly analogous to the human Achilles tendon in its capacity for energy storage, injury profile and extent of specialization(Biewener, 1998; Patterson-Kane & Rich, 2014). Availability of samples enabled us to explore the extensive adaptation processes associated with late stage development, contrasting paired positional and energy-storing equine tendons through pre- and post-natal development.

Using this model, we investigate the process and drivers of functional adaptation, when tendons transition from an absence of loading (foetal: mid to end (6 to 9 months) gestation, and 0 days: full-term foetuses, and foals that did not weight-bear); through to weight-bearing (0-1 month) and then to an increase in body weight and physical activity (3-6 months; and 1-2 years). We hypothesise that early in development during gestation, the fibre and matrix phases of functionally distinct tendons have identical compositional profiles and mechanical properties, with tissue specialisation occurring as an adaptive response to the mechanical stimulus of load-bearing, predominantly in the matrix phase of the elastic energy-storing tendon.

## Results

### Mechanical adaptation is localised to the matrix phase

First, we determined how the mechanical properties of the fibre and matrix phase develop in tendon, with a particular focus on the temporospatial nature of mechanical adaptation to functional specialisation. Individual fascicles were dissected and tested as fibre samples, while an isolated region of interfascicular matrix, matrix phase, was tested by shearing fascicles apart. Samples were subjected to preconditioning followed by a pull to failure (Supplementary Figure 1). The yield point of samples was identified, denoting the point at which the sample became irreversibly damaged and was unable to recover from the applied load, and the sample failure properties also recorded, highlighting the maximum stress and strain the sample could withstand.

A significant increase in fibre phase yield and failure properties was evident when comparing embryonic fibre samples to those acquired immediately at birth. However, data indicate minimal distinction in fibre phase mechanics between functionally distinct tendons (Figure 1) and, significantly, no specialisation for energy storage in response to loading during postnatal development.

**Figure 1.**
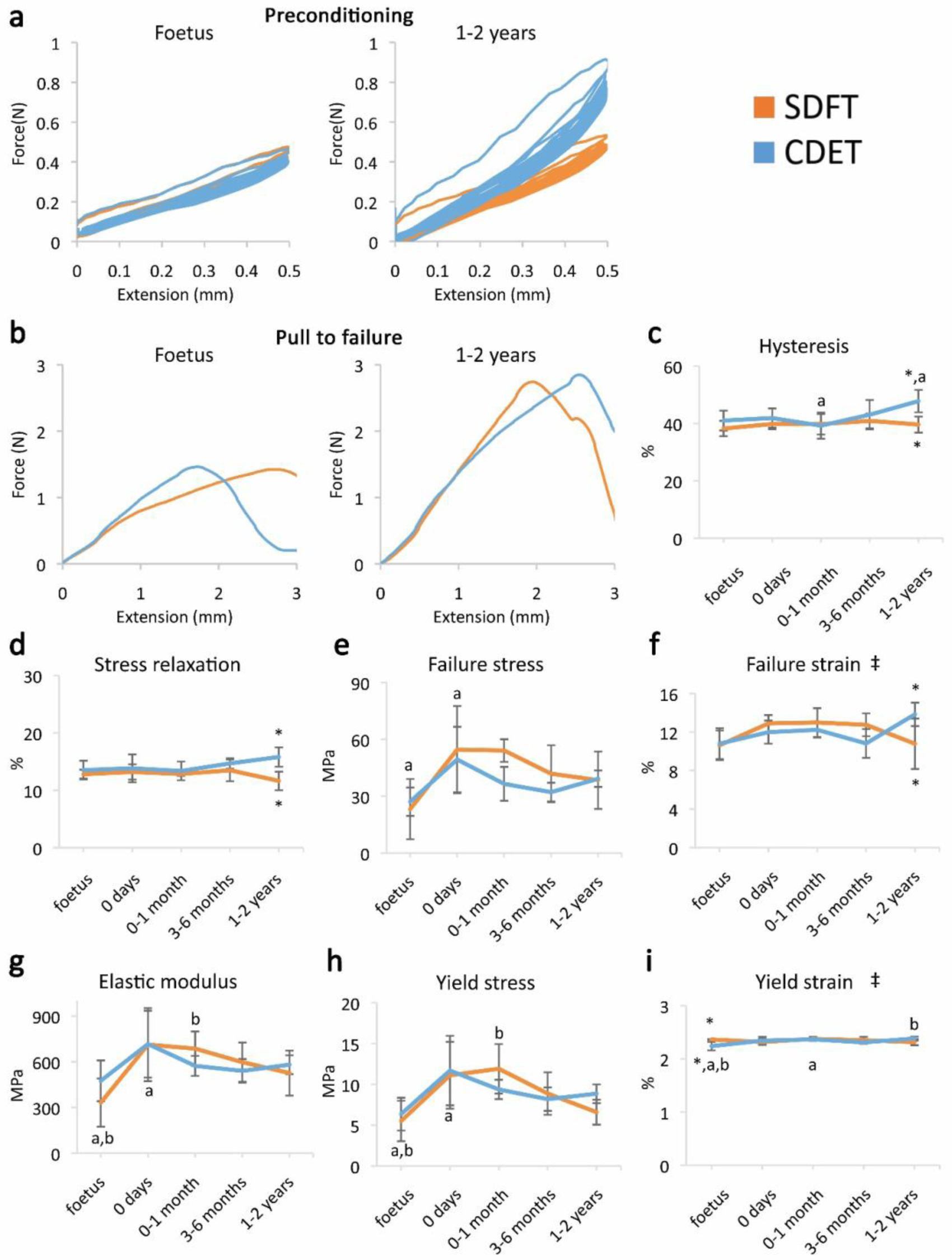
Fibre phase response to mechanical testing shows increase in strength with development but few significant differences between tendon types, indicating that the fibre phase shows minimal structural specialisation in response to loading. (**a**) Representative curves for 10 preconditioning cycles for the SDFT and CDET fibre phase in the foetus and 1-2 years age group. (**b**) Representative force-extension curves to failure for the SDFT and CDET fibre phase in the same age groups. (**c-i**) Mean SDFT and CDET fibre phase biomechanical properties are presented across development, with data grouped into age groups: foetus, 0 days (did not weight-bear), 0-1 month, 3-6 months, 1-2 years. ‡ significant interaction between tendon type and development, * significant difference between tendons, a-b significant difference between age groups. Error bars depict standard deviation.

Contrasting with fibre phase mechanics, the failure properties of the matrix phase continued to alter throughout development with failure properties increasing markedly from 6 months onwards. We also identify the emergence of an extended region of low stiffness at the start of the loading curve (ie an extended toe region) specific to the SDFT matrix phase pull to failure curve. This indicates less resistance to extension, and together with the concomitant increase in matrix yield force and extension at yield, demonstrates development of an overall greater capacity for extension in the SDFT matrix phase behaviour. A summary of these findings is achieved by plotting the amount of matrix phase extension at different percentages of failure force (Figure 2b,h-k), highlighting how the matrix phase of the energy-storing SDFT became significantly less stiff than that of the positional CDET during the initial toe region of the loading curve as the tendon adapts.

**Figure 2.**
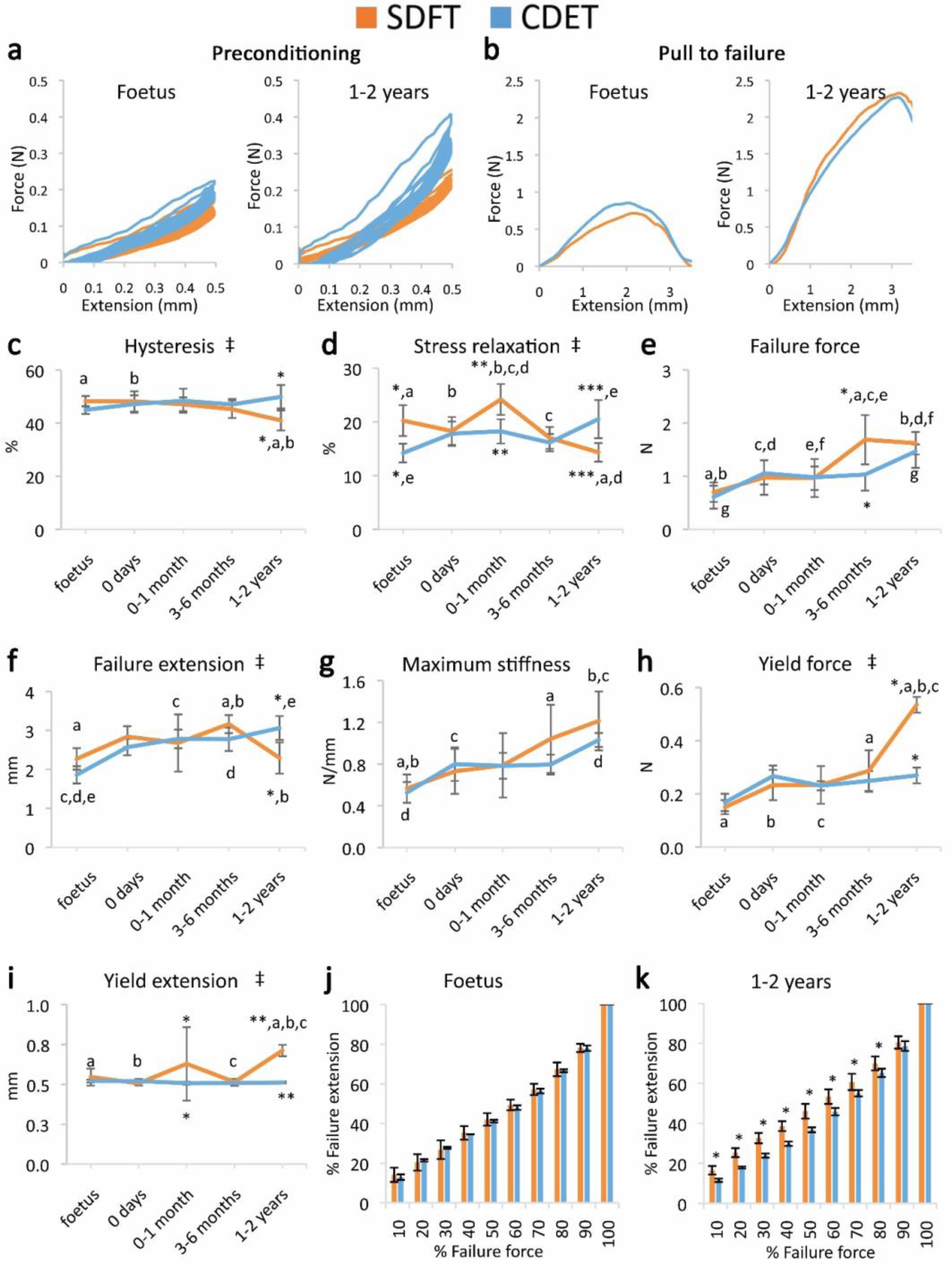
Mechanical testing of the matrix phase shows an equivalent increase in failure properties between the SDFT and CDET with development, but development of an extended low stiffness toe region and more elastic behaviour in the SDFT. (**a**) Representative curves for 10 preconditioning cycles for the SDFT and CDET matrix phase in the foetus and 1-2 years age group. (**b**) Representative force-extension curves to failure for the SDFT and CDET matrix phase in the same age groups. (**c-i**) Mean SDFT and CDET matrix phase biomechanical properties are presented across development, with data grouped into age groups: foetus, 0 days (did not weight-bear), 0-1 month, 3-6 months, 1-2 years. (**j-k**) To visualise the extended low stiffness toe region in the SDFT matrix phase, the amount of matrix phase extension at increasing percentages of failure force is presented, comparing the SDFT and CDET in the foetus and 1-2 years age group. ‡ significant interaction between tendon type and development, * significant difference between tendons, a-g significant difference between age groups. Error bars depict standard deviation.

The viscoelastic properties of the developing matrix phase also showed significant interactions between tendon type and development, with matrix viscoelasticity significantly decreasing with development specifically in the energy-storing SDFT (Figure 2a,c-d), resulting in specialisation towards a more energy efficient structure.

### Structural adaptation is localised to the matrix phase

Having described mechanical adaptation of the matrix phase to meet functional demand, we next performed a histological and immunohistochemical comparison of developing energy-storing and positional tendons to determine how temporospatial structural adaptation may dictate this evolving mechanical behaviour.

The energy-storing SDFT and positional CDET appeared histologically similar in the foetus, in both instances showing surprisingly poor demarcation of the matrix phase, which only became structurally distinct after birth and the initiation of loading (Figure 3a). Fibre development was generally consistent in both tendon types with cellularity and crimp showing a reduction with development, cells displaying more elongated nuclei, and collagen showing a more linear organisation. In contrast, the matrix phase demonstrated divergence between tendon types with only the SDFT matrix phase showing an increase in cellularity following tendon loading and a retention of matrix phase width throughout development (Figure 3, Supplementary Figure 2).

**Figure 3.**
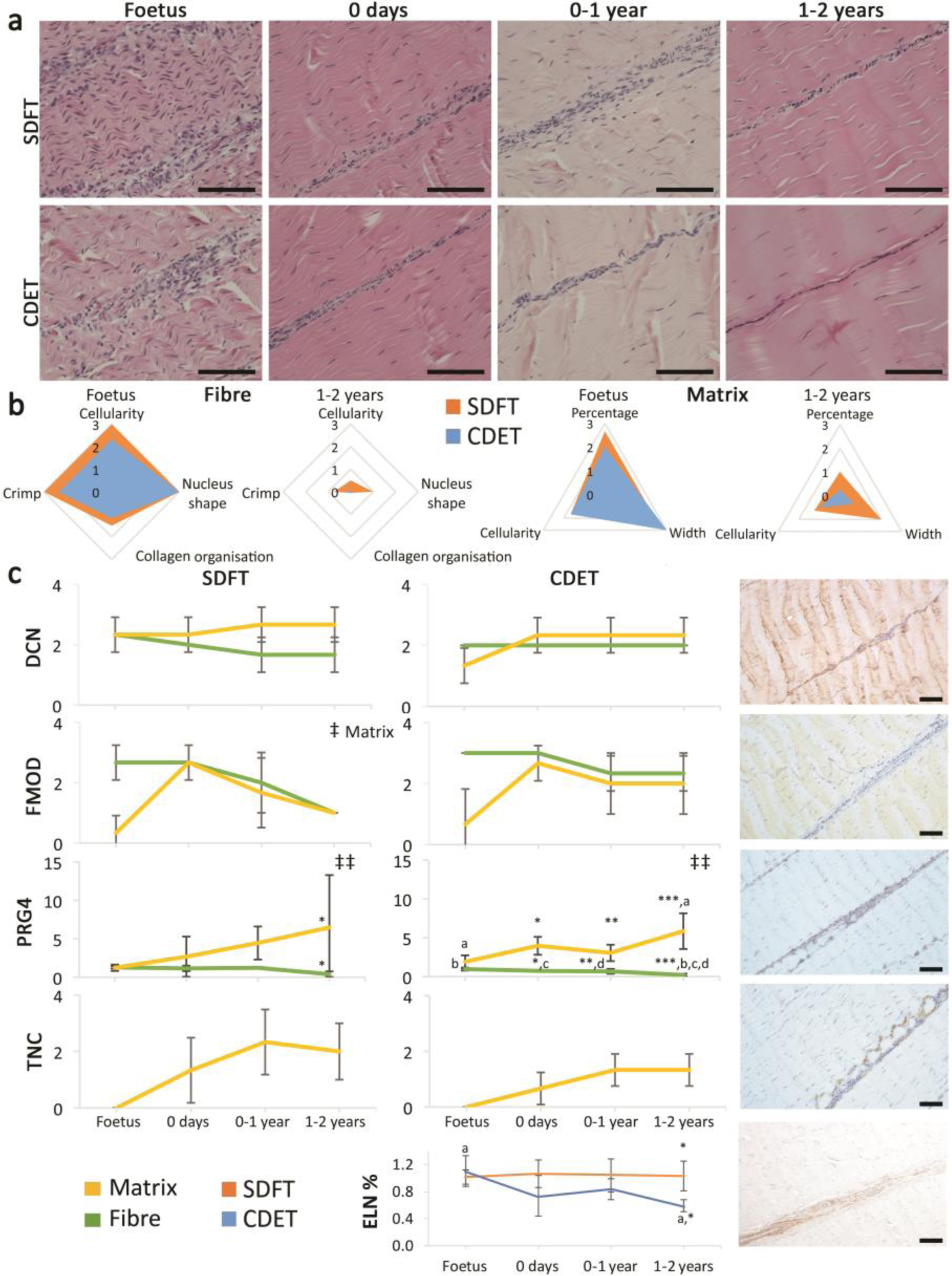
The SDFT and CDET are histologically similar at birth and differentiate with development especially in the matrix phase. (**a**) Representative images of H&E sections of the SDFT and CDET demonstrate structural development: foetus, 0 days (did not weight-bear), 0-1 year, and 1-2 years whilst (**b**) Radar plots enable the mean histology scores of the fibre and matrix phase for the SDFT and CDET to be compared between the foetus and 1-2 years age group (all data shown in Supplementary Figure 2). A decrease in cell numbers, crimp, and matrix phase width is visible with progression of age, and the aspect ratio of cells in the fibre phase increases. (**c**) Use of immunohistochemical assays shows divergence of PGR4 (lubricin) and elastin with maturation between functionally distinct tendons. Matrix and fibre phase staining scores are shown for decorin (DCN), fibromodulin (FMOD), lubricin (PRG4), and tenascin-C (TNC) in the SDFT and CDET, alongside representative images of immunohistochemical staining in the postnatal SDFT. A quantitative measure of elastin (ELN) is provided as percentage of wet weight, alongside a representative image of immunohistochemical staining in the postnatal SDFT. Staining scores for elastin are provided in Supplementary Figure 3. ‡ significant change in tendon phase with development, ‡‡ significant interaction between tendon phase and development, * significant difference between tendons, a-d significant difference between age groups. Scale bar 100 µm. Error bars depict standard deviation.

The abundance of major ECM proteins was also generally consistent across fibre and matrix regions in the foetus, with divergence of protein composition between phases only evident with further development. Notably adaptation was driven by changes in non-collagenous ECM components specifically, levels of which reduced in the fibre phase and increased dramatically in the matrix phase through postnatal development (Figure 3c, Supplementary Figure 3). Of particular note, we demonstrate the distribution of PRG4 (commonly known as lubricin) and TNC predominantly in the matrix phase of tendons. We also demonstrate that elastin is preferentially localised to the matrix phase with its abundance decreasing only in the CDET with development. Furthermore, we show how structural changes do not manifest until after birth and the initiation of loading, but that structural adaptation to loading then occurs over a period of months, involving both upregulation and downregulation of different ECM constituents.

### Adaptation relies on evolution of matrix phase composition only

To explore these concepts in further detail and to scrutinise the capacity for ECM adaptation, proteomic methodologies were adopted. With the mechanical and histological data identifying that functional adaptation is particular to the energy-storing SDFT, mass spectrometry analysis focused on a more detailed comparison of the matrix and fibre phase development and adaptation in this tendon specifically.

Our results demonstrated that the proteomic profile of the matrix phase was more complex (more identified proteins) and a higher percentage of matrix phase proteins were cellular, supporting the histological findings of a more cellular matrix phase (Supplementary Figure 4). Notably, despite the two phases being structurally distinct, they had 14 collagens and 11 proteoglycans in common (Supplementary Table 4). Overall, proteomic heatmap analysis correlated very strongly with immunohistochemical findings, showing that alterations in the fibre phase proteome reduced through development, with minimal changes following the initiation of loading (Table 1 and 2, Figure 4a), whilst numerous matrisome and matrisome-related proteins progressively increase in abundance through development in the matrix phase (Table 1 and 2). Detailed consideration of protein changes also highlights that post loading changes in the matrix phase appear more specific to proteoglycans and glycoproteins.

**Table 1.**
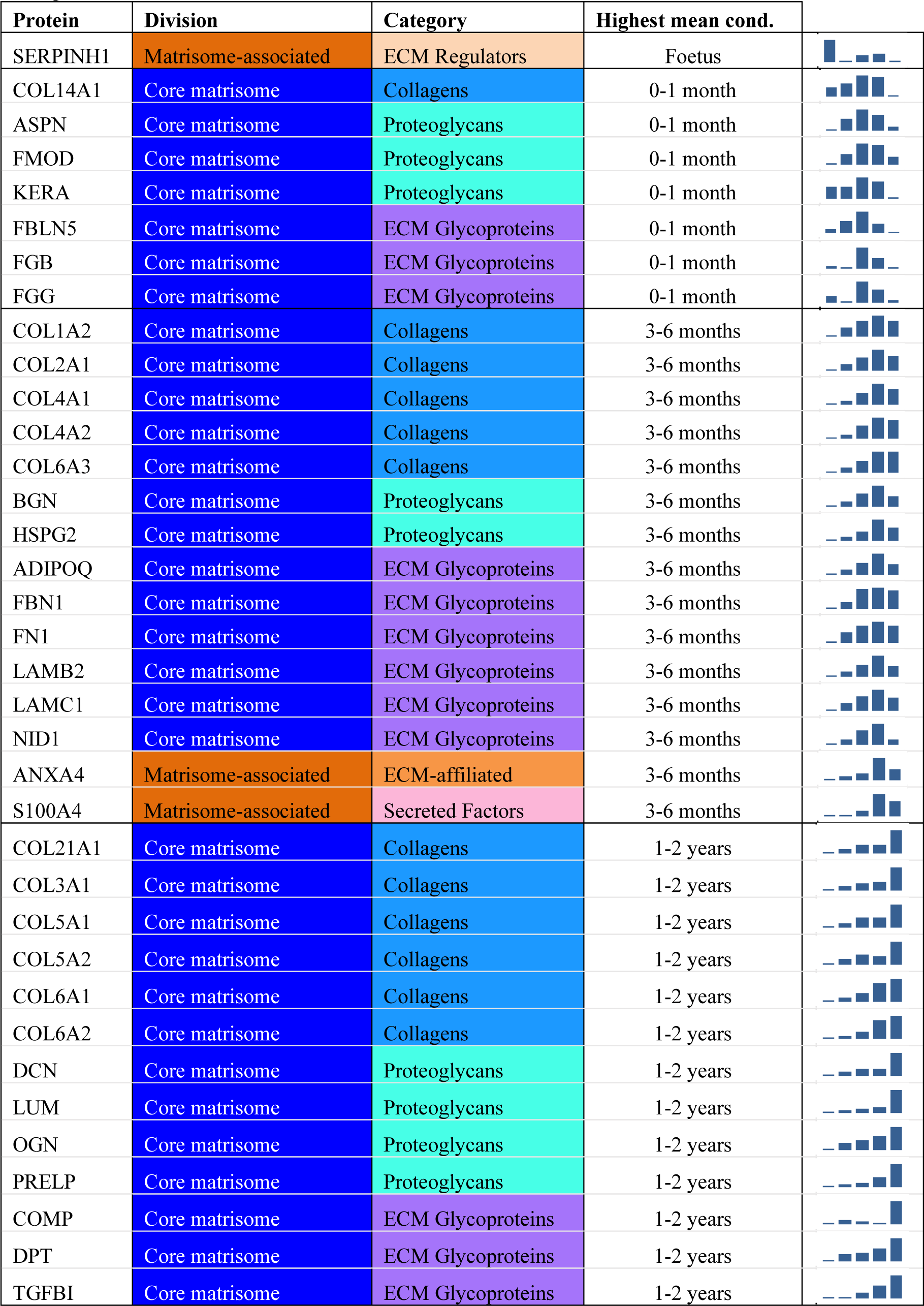
Matrix phase differentially abundant matrisome and matrisome-associated proteins through development organised by highest mean condition (*p*<0.05, fold change≥2). Proteins are arranged into colour coded divisions and categories. Bar graphs profile the relative abundance of each protein at each development stage, with the development age reporting the highest mean protein level also specified.

| Protein | Division | Category | Highest mean cond. |
| --- | --- | --- | --- |
| SERPINH1 | Matrisome-associated | ECM Regulators | Foetus |
| COL14A1 | Core matrisome | Collagens | 0-1 month |
| ASPN | Core matrisome | Proteoglycans | 0-1 month |
| FMOD | Core matrisome | Proteoglycans | 0-1 month |
| KERA | Core matrisome | Proteoglycans | 0-1 month |
| FBLN5 | Core matrisome | ECM Glycoproteins | 0-1 month |
| FGB | Core matrisome | ECM Glycoproteins | 0-1 month |
| FGG | Core matrisome | ECM Glycoproteins | 0-1 month |
| COL1A2 | Core matrisome | Collagens | 3-6 months |
| COL2A1 | Core matrisome | Collagens | 3-6 months |
| COL4A1 | Core matrisome | Collagens | 3-6 months |
| COL4A2 | Core matrisome | Collagens | 3-6 months |
| COL6A3 | Core matrisome | Collagens | 3-6 months |
| BGN | Core matrisome | Proteoglycans | 3-6 months |
| HSPG2 | Core matrisome | Proteoglycans | 3-6 months |
| ADIPOQ | Core matrisome | ECM Glycoproteins | 3-6 months |
| FBN1 | Core matrisome | ECM Glycoproteins | 3-6 months |
| FN1 | Core matrisome | ECM Glycoproteins | 3-6 months |
| LAMB2 | Core matrisome | ECM Glycoproteins | 3-6 months |
| LAMC1 | Core matrisome | ECM Glycoproteins | 3-6 months |
| NID1 | Core matrisome | ECM Glycoproteins | 3-6 months |
| ANXA4 | Matrisome-associated | ECM-affiliated | 3-6 months |
| S100A4 | Matrisome-associated | Secreted Factors | 3-6 months |
| COL21A1 | Core matrisome | Collagens | 1-2 years |
| COL3A1 | Core matrisome | Collagens | 1-2 years |
| COL5A1 | Core matrisome | Collagens | 1-2 years |
| COL5A2 | Core matrisome | Collagens | 1-2 years |
| COL6A1 | Core matrisome | Collagens | 1-2 years |
| COL6A2 | Core matrisome | Collagens | 1-2 years |
| DCN | Core matrisome | Proteoglycans | 1-2 years |
| LUM | Core matrisome | Proteoglycans | 1-2 years |
| OGN | Core matrisome | Proteoglycans | 1-2 years |
| PRELP | Core matrisome | Proteoglycans | 1-2 years |
| COMP | Core matrisome | ECM Glycoproteins | 1-2 years |
| DPT | Core matrisome | ECM Glycoproteins | 1-2 years |
| TGFBI | Core matrisome | ECM Glycoproteins | 1-2 years |

**Table 2.**
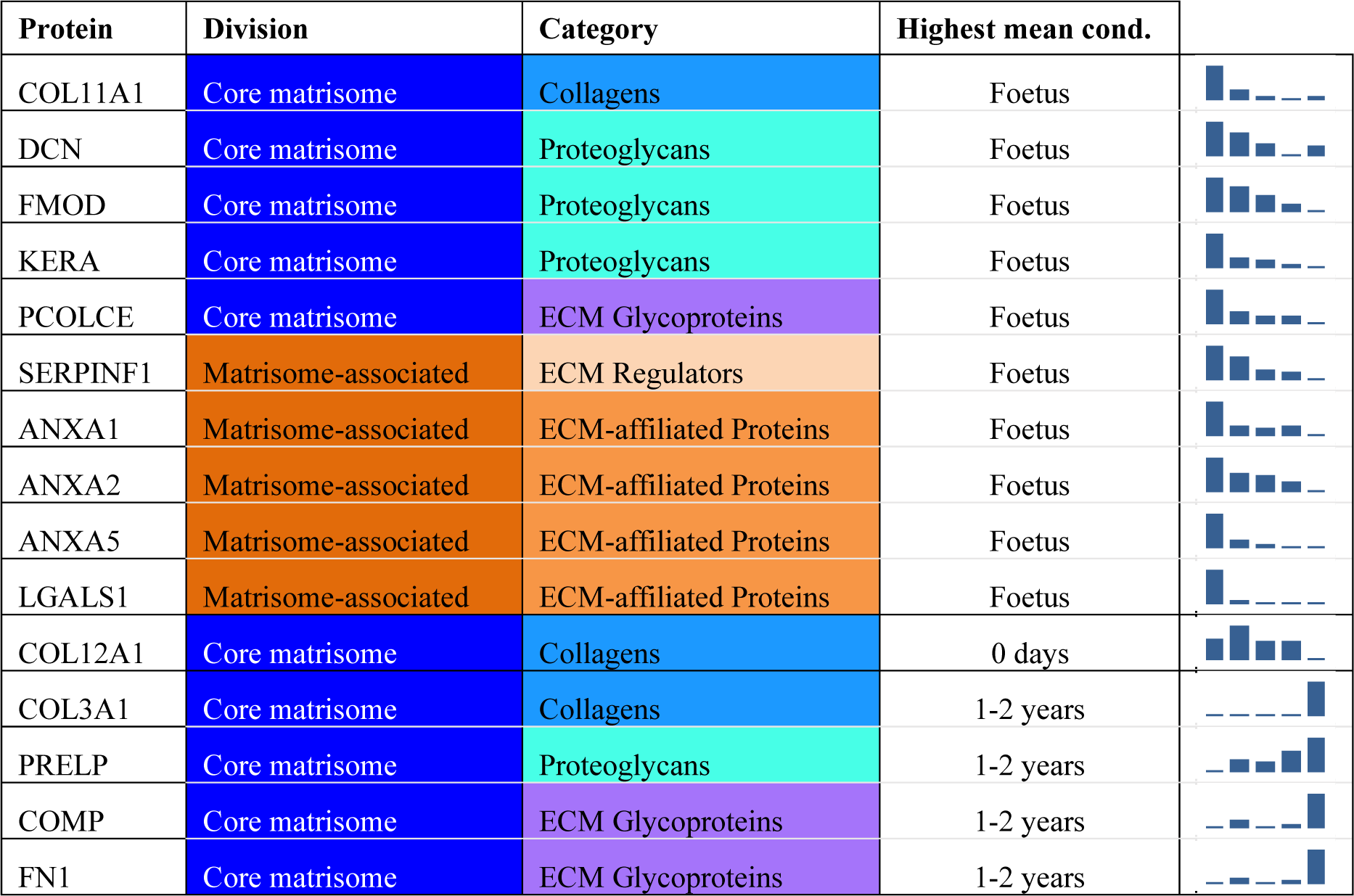
Fibre phase differentially abundant matrisome and matrisome-associated proteins through development organised by highest mean condition (*p*<0.05, fold change≥2). Proteins are arranged into colour coded divisions and categories. Bar graphs on the right profile the relative abundance of each protein at each development stage, with the development age reporting the highest mean protein level also specified.

| Protein | Division | Category | Highest mean cond. |
| --- | --- | --- | --- |
| COL11A1 | Core matrisome | Collagens | Foetus |
| DCN | Core matrisome | Proteoglycans | Foetus |
| FMOD | Core matrisome | Proteoglycans | Foetus |
| KERA | Core matrisome | Proteoglycans | Foetus |
| PCOLCE | Core matrisome | ECM Glycoproteins | Foetus |
| SERPINF1 | Matrisome-associated | ECM Regulators | Foetus |
| ANXA1 | Matrisome-associated | ECM-affiliated Proteins | Foetus |
| ANXA2 | Matrisome-associated | ECM-affiliated Proteins | Foetus |
| ANXA5 | Matrisome-associated | ECM-affiliated Proteins | Foetus |
| LGALS1 | Matrisome-associated | ECM-affiliated Proteins | Foetus |
| COL12A1 | Core matrisome | Collagens | 0 days |
| COL3A1 | Core matrisome | Collagens | 1-2 years |
| PRELP | Core matrisome | Proteoglycans | 1-2 years |
| COMP | Core matrisome | ECM Glycoproteins | 1-2 years |
| FN1 | Core matrisome | ECM Glycoproteins | 1-2 years |

**Figure 4.**
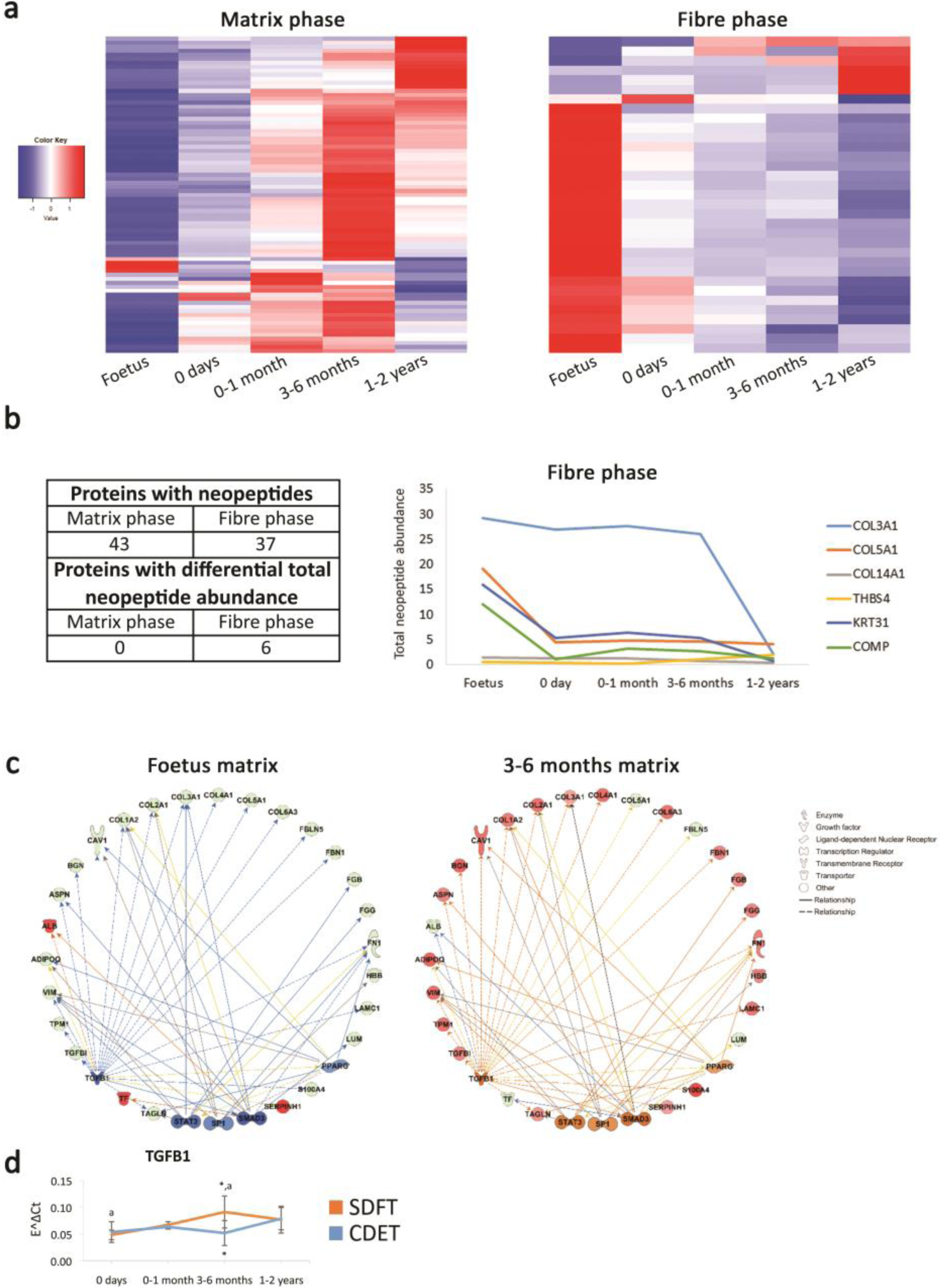
The fibre phase proteome remains the same during postnatal development and tendon loading whereas the matrix phase starts changing following tendon loading in postnatal development. (**a**) Heatmap of differentially abundant proteins in foetus, 0 days (did not weight-bear), 0-1 month, 3-6 months, and 1-2 years SDFT matrix and fibre phases (*p*<0.05, fold change ≥2). Heatmap colour scale ranges from blue to white to red with blue representing lower abundance and red higher abundance. (**b**) Proteins with identified neopeptides and proteins showing differential total neopeptide abundance across age groups. Graph of proteins showing differential total neopeptide abundance in the SDFT fibre phase across development (*p*<0.05, fold change≥2, FDR 5%). No proteins showed differential total neopeptide abundance in the matrix phase. (**c**) IPA networks for TGFB1 in the foetus and at 3-6 months in the SDFT matrix phase. TGFB1 regulation in the matrix phase is predicted to be inhibited in the foetus and activated at 3-6 months in the SDFT. Red nodes, upregulated proteins, green nodes, downregulated proteins, intensity of colour is related to higher fold-change, orange nodes, predicted upregulated proteins in the dataset, blue nodes, predicted downregulated proteins. (**d**) Whole tendon relative mRNA expression for TGFB1 in the SDFT and CDET during postnatal development. * significant difference between tendons, a significant difference between age groups. Error bars depict standard deviation.

Proteomic data also enable insight into the turnover of proteins in the different tendon phases, through a comparison of the neopeptides produced by protein breakdown(Chavaunne T. Thorpe, Peffers, et al., 2016). In the current study, we are able to profile the temporal nature of fibre and matrix turnover, demonstrating that both phases display turnover during development but that fibre phase turnover slows down towards the end of maturation, whilst matrix phase turnover rates are maintained, suggesting structural and/or compositional plasticity (Figure 4b).

Having identified the matrix phase as the location of functional adaptation, we next investigated the regulation of this process, to detect targets for modulation in regeneration strategies addressing functional impairment. Pathway analysis of the differentially abundant proteins across age groups supported an ECM-integrin-cytoskeleton to nucleus signalling pathway for the mediation of the observed mitogenesis and matrigenesis in response to tendon loading. In addition, TGF-β1 was highlighted as a regulator of ECM organisation and functional adaptation, upregulated in the energy-storing tendon following loading (Figure 4).

## Discussion

In this study, we describe the process and drivers of functional adaptation in tendon development integrating mechanical, structural, and compositional analysis in tendons transitioning from an absence of loading through to weight-bearing and then to an increase in body weight and physical activity. Whilst the limited previously available data on the development of tendon gross mechanical properties show an increase in mechanical properties with development(Ansorge et al., 2011; Cherdchutham, Meershoek, et al., 2001), no such phase-specific analysis of the development of tissue mechanics has been carried out previously. Similarly, available research into tendon morphogenesis and maturation has previously focused on the development of the collagenous network that comprises the fibrous phase of the tendon and is often focussed on early foetal development(Kalson et al., 2011; Marturano et al., 2013; Pan et al., 2018).

Here, examining the fibre and matrix phase mechanical properties independently, we show minimal distinction in fibre phase mechanics between functionally distinct tendons and, significantly, no specialisation for energy storage in response to loading during postnatal development. In contrast, the matrix phase mechanical properties display continuous alterations through development with properties in the foetus being comparable between functionally distinct tendons, and a low stiffness region emerging in the initial non-linear region (toe region) of the pull to failure curve of the SDFT matrix phase only following tendon loading postnatally. This is coupled with a concomitant increase in matrix force and extension at yield, point at which the sample became irreversibly damaged, highlighting that the energy-storing SDFT matrix phase becomes significantly less stiff than that of the positional CDET during the initial toe region of the loading curve. We have previously indicated that this low stiffness behaviour allows sliding between the fascicles of the fibre phase, enabling non-uniform loading of tissue and is fundamental for effective extension and recoil in energy-storing tendons(Chavaunne T. Thorpe et al., 2015). Furthermore, with ageing the low stiffness behaviour of the matrix phase of energy-storing tendons is lost, possibly contributing to disease development(Chavaunne T. Thorpe et al., 2013). The only other studies considering the mechanical properties of developing tendons have focused simply on changes in whole tissue mechanics, and have thus not been able to identify the drivers of change within the tissue(Ansorge et al., 2011; Cherdchutham, Meershoek, et al., 2001). Here, we identify that the matrix phase is the key region in which mechanical adaptation to meet function occurs, and that this occurs after the initiation of loading, primarily 1-2 years postnatally.

We subsequently examine how temporospatial structural adaptation may dictate this evolving mechanical behaviour and uncover a divergence in structural characteristics between tendon types in the matrix phase only, with a retention of matrix phase width throughout development and an increase in cellularity in the SDFT matrix phase only. It is well recognised that foetal and early postnatal tendons are highly cellular, and cellularity is generally considered to decrease postnatally(Russo et al., 2015; Stanley et al., 2008), but here we show that the described reduction in cellularity only occurs in the fibre phase, and cellularity in fact increases in the matrix phase with development. By following the alterations in cellularity across fibre and matrix, here we can determine that the marked difference in regional cellularity is likely driven by a maintenance of cell numbers following tendon loading in the thinning matrix region, while cell numbers in the fibre region appear to reduce as a result of the fibre phase ECM increase. Greater cellularity is commonly associated with a requirement for rapid adaptive organisation of ECM components(Russo et al., 2015), suggesting the matrix phase, particularly in the energy-storing tendons, may adapt to be more mechanoresponsive, a necessary aspect of healthy homeostatic maintenance of a tissue. Immunohistochemical analysis reveals the distribution of major ECM proteins is consistent across tendons in the foetus and becomes distinct across phases through development. Of note, PRG4 (lubricin), a large proteoglycan which is important in ECM lubrication, is found mainly distributed in the matrix phase of tendons. Using a lubricin-knockout mouse, this proteoglycan has been demonstrated to facilitate interfascicular sliding(Kohrs et al., 2011), indicating that this structural adaptation may be key in achieving the previously identified mechanical adaptation in the energy-storing tendon matrix phase. In addition, Kostrominova and Brooks (2013)(Kostrominova & Brooks, 2013) report PRG4 expression as wells as elastin expression decreased with ageing in rat tendon suggesting an association with an increased risk of disease with ageing. We also demonstrate that elastin is preferentially localised to the matrix phase, potentially having a role in the capacity for matrix recoil after loading which is necessary for the healthy function of energy-storing tendons(Godinho et al., 2017; Ritty et al., 2002). Further supporting a role for elastin in the energy-storing function, elastin appears to be redundant in positional tendons with its abundance decreasing in the CDET with development. Together, these findings show that structural adaptation of tendon post loading is primarily focused to the matrix phase, and observed predominantly in the energy-storing SDFT.

Compositional analysis of the energy-storing SDFT phases using mass spectrometry corroborates the immunohistochemical analysis and shows a more complex proteomic profile for the matrix phase. It additionally shows that abundance of the majority of proteins in the fibre phase is higher in the foetus and reduces through development with minimal changes following the initiation of loading, whereas in the matrix phase numerous ECM and ECM-related proteins progressively increase in abundance following tendon loading and through development. Neopeptide analysis demonstrated fibre phase ECM protein turnover slows down towards the end of maturation, whilst matrix phase ECM protein turnover rates are maintained. Once a tendon is mature, little collagen turnover occurs(Birch, 2007; KM et al., 2013) and we have previously shown that the minimal turnover in mature tendon is focused to the matrix phase(C. T. Thorpe et al., 2015; Chavaunne T. Thorpe et al., 2010; Chavaunne T. Thorpe, Peffers, et al., 2016). The maintenance of turnover rates observed in the matrix phase here suggests structural and/or compositional plasticity of the matrix phase. Integrated, our data convincingly show a continual temporal change in the matrix phase proteome specifically, which would enable adaptation and specialisation to the load environment, and highlight the compositional plasticity of the matrix phase in responding to dynamic altered conditions such as those occurring during development and regeneration. It is critical that the difference in capacity for functional adaptation across fibre and matrix phases identified here is considered if regenerative medicine and tissue engineering approaches are to be successful. Here, we demonstrate the temporal pattern of structure-function adaptation, with compositional changes occurring in the first months after loading, and leading to the mechanical specialisation we have previously observed in adult energy-storing tendon(Birch, 2007; C T Thorpe et al., 2012; Chavaunne T. Thorpe et al., 2015). With the fibrous phase primarily responsible for the mechanical strength of a tissue, biomaterial and regenerative medicine studies have unsurprisingly placed considerable emphasis on this region to date(Sensini et al., 2019; Watts et al., 2017). Here, we not only highlight the importance of the matrix phase in modulating mechanical behaviour, but also demonstrate how the matrix phase must be targeted to support adaptation and optimum tissue quality.

Finally, pathway analysis of our proteomic data highlighted TGF-β1 as a regulator of ECM organisation and functional adaptation, predicted to be upregulated following loading in the energy-storing tendon. TGF-β is known to have a role in cellular mechanobiology and connective tissue homeostasis, regulating ECM synthesis and remodelling in a force-dependent way following mechanical stimulation, to specify the quality of the ECM and help coordinate cytoskeletal tension(Maeda et al., 2011). A previous study of developing chick tendon detected TGF-β1 staining in the matrix phase only during development, highlighting its localised distribution in development(Kuo et al., 2008). In the current study, we are also able to associate TGF-β expression with functional adaptation of the matrix phase of tendon. In addition to tissue development and homeostasis, TGF-β1 is involved in connective tissue injury and repair with abnormal expression levels reported in both processes suggesting a pleiotropic mode of action(Gao et al., 2019). The above may suggest a role for TGF-β1 in tissue development and homeostasis and that its dysregulation is associated with tissue injury and repair expression levels.

### Outlook

We demonstrate for the first time that functional adaptation in tendon is predominantly reliant on adaptation of the metabolically active matrix phase, which responds to the mechanical environment through TGF-β signalling, resulting in modulations in ECM turnover and composition to fine-tune mechanical properties. Traditionally, the matrix phase of connective tissues has received considerably less attention than the fibre phase, with regenerative medicine, biomimetics and biomechanics studies all largely focused on investigating and recapitulating the organisation and mechanical properties of the collagenous fibrous network. Following tendon injury, normal tissue architecture is not recovered, and in particular, the cellular matrix phase is not regenerated. There is great potential gain from understanding the convergence of biology underpinning adaptation, function and pathology and here, we propose a paradigm shift to consider the metabolically active matrix phase as a key target for regenerative medicine strategies aimed at addressing functional impairment of tendons and other connective tissues following disease. Regeneration of the matrix phase following tendon injury could be key for tendon health and low re-injury risk.

## Materials and Methods

### Experimental design

Using an equine tendon model, we investigate the process and drivers of functional adaptation in the SDFT and CDET, two functionally distinct tendons, when tendons transition from an absence of loading (foetal: mid to end (6 to 9 months) gestation, and 0 days: full-term foetuses, and foals that did not weight-bear); through to weight-bearing (0-1 month) and then to an increase in body weight and physical activity (3-6 months; and 1-2 years). We use a phase-specific approach to characterise each tendon phase independently, by comparing fascicles (fibre phase) and inter-fascicular matrix (IFM; matrix phase) mechanical properties, structure and composition.

For this purpose, we used mechanical testing, histological and immunohistochemical analysis and mass spectrometry analysis following laser capture microdissection. Sample size was selected based on previous experiments and restricted by sample availability and the cost of mass spectrometry analysis.

### Sample collection

Both forelimbs were collected from foetuses and foals aged 0-2 years (n=19) euthanised for reasons unrelated to this project at a commercial abattoir or equine practices following owner consent under ethical approval for use of the cadaveric material granted by the Veterinary Research Ethics Committee, School of Veterinary Science, University of Liverpool (VREC352). Collected tendons were split in the following age groups: Foetus (between 6 and 9 months of gestation; n=4); 0 days (full-term foetuses (average gestation 11-12 months) and foals that did not weight-bear; n=4): 0-1 month (n=3); 3-6 months (n=4); 1-2 years (n=4). The SDFT and CDET from one forelimb were dissected and wrapped in phosphate-buffered saline dampened tissue paper and foil and stored at −80 °C for biomechanical testing. Two 1-2 cm segments from the mid-metacarpal area of the SDFT and CDET of the other forelimb were dissected, and one fixed in 4% paraformaldehyde for histology and immunohistochemistry, and the other snap frozen in isopentane and stored at −80 °C for laser capture microdissection.

### Biomechanical testing of the fibre phase

On the day of testing, samples were defrosted within their tissue paper wrap, then immediately prepared for testing. Fascicles (fibre phase samples) were dissected from the mid-metacarpal region of the SDFT and CDET and subjected to a quasi-static test to failure according to Thorpe et al. (2015)(Chavaunne T. Thorpe et al., 2015). Briefly, prior to testing, the diameter of each fascicle was measured along a 1 cm length in the middle of the fascicle with a non-contact laser micrometre (LSM-501, Mitotuyo, Japan, resolution = 0.5 um) and the smallest diameter recorded and used to calculate cross-sectional area (CSA), assuming a circular shape.

Fibre phase samples were loaded in an electrodynamic testing machine (Instron ElectroPuls 1000) equipped with a 250 N load cell and pneumatic grips (4 bar) coated with rubber and sandpaper to prevent sample slippage(C T Thorpe et al., 2012). The distance between the grips was set to 20 mm and fibre phases preloaded to 0.1 N (approx. 2% fibre phase failure load) to remove any slack in the sample. Following preload, the distance between the grips was recorded as the gauge length, then fibre phase samples preconditioned with 10 loading cycles between 0 and 3% strain (approximately 25% failure strain) using a sine wave at 1 Hz frequency. Immediately after preconditioning, fibre samples were pulled to failure at a strain rate of 5%/s. Force and extension data were continuously recorded at 100 Hz during both preconditioning and the quasi-static test to failure. Acquired data were smoothed to reduce noise before calculations with a 3_rd_ order Savitzky-Golay low pass filter, with a frame of 15 for the preconditioning data and 51 for the pull to failure data.

Using the preconditioning data, the total percentage hysteresis and stress relaxation were calculated, between the first and last preconditioning cycle. Failure force, extension, stress, and strain were calculated from the test to failure, and a continuous modulus calculated across every 10 data points of each stress strain curve, from which the maximum modulus value was determined. The point of maximum modulus was defined as the yield point from which yield stress and yield strain were determined.

### Biomechanical testing of the matrix phase

On the day of testing, tendons were defrosted within their tissue paper wrap, and matrix phase samples immediately dissected and prepared for biomechanical testing as described previously by Thorpe et al. (2012)(C T Thorpe et al., 2012). Briefly, a group of two adjacent intact fascicles (bound by the matrix phase) were dissected, after which the opposing end of each fascicle was cut transversely 10 mm apart, to leave a consistent 10 mm length of intact matrix phase that could be tested in shear (Supplementary Figure 1).

Utilising the same electrodynamic testing machine and pneumatic grips as described for the fibre phase, the intact end of each fascicle was gripped with a grip to grip distance of 20 mm, and a pre-load of 0.02 N (approx. 1% matrix phase failure load) applied. Matrix phase samples were preconditioned with 10 cycles between 0 and 0.5 mm extension (approx. 25% failure extension) using a sine wave at 1 Hz frequency, then pulled to failure at a speed of 1 mm/s. Force and extension data were continuously recorded at 100 Hz during both preconditioning and the quasi-static test to failure. Acquired data was smoothed to reduce noise before calculations with a 3_rd_ order Savitzky-Golay low pass filter, with a frame of 15 for the preconditioning data and 51 for the pull to failure data.

Total percentage hysteresis and stress relaxation were again calculated between the first and last preconditioning cycle. Failure force and extension were determined from the quasi-static pull to failure curve, and a continual stiffness curve was calculated across every 10 data points of the curve, from which maximum stiffness was determined, and yield force and yield extension at maximum stiffness reported. Based on previous data demonstrating notable differences in the toe region of the matrix phase curve of functionally distinct tendons, the shape of failure curves was also compared between samples by calculating the amount of matrix phase extension at different percentages of matrix phase failure load(Chavaunne T. Thorpe et al., 2015).

### Histology scoring

Paraformaldehyde-fixed paraffin-embedded longitudinal SDFT and CDET segments were sectioned at 6 µm thickness and stained with H&E for histologic examination and scoring (n=13; 3 from each age group, 4 from 1-2 years age group). The examined histologic variables are reported in Supplementary Table 1.

For parameters scored by investigators, the sections were blinded and histologic variables assigned a grade from 0 to 3 by two independent investigators. Weighted Kappa showed moderate to good agreement in all instances, hence the average of the two scores was used. Other histologic variables were measured using image analysis (TissueFaxs, Tissuegnostics and Adobe Photoshop CS3) and then assigned a grade from 0 to 3. Cumulative scores for the fibre and matrix phase for each horse were obtained by summing the scores of the fibre and matrix phase variables, respectively, excluding matrix phase percentage to ensure matrix phase dimensions were not over-weighted in final reporting.

### Immunohistochemistry

Immunohistochemical analysis for DCN, FMOD, PRG4, TNC and ELN was carried out on paraformaldehyde-fixed paraffin-embedded longitudinal SDFT and CDET sections (6 µm thickness) (n=12; 3 from each age group) as previously described by Zamboulis et al. (2013)(Zamboulis et al., 2013). Antigen retrieval was carried out with 0.2 U/mL Chondroitinase ABC (C2905, Sigma, Merck, Darmstadt, Germany) at 37 °C for two hours for DCN, FMOD, PRG4, and TNC or with 4800 U/mL hyaluronidase (H3506, Sigma, Merck, Darmstadt, Germany) at 37 °C for two hours for ELN. Primary antibodies were used at a concentration of 1:1500 for DCN (mouse IgG), 1:400 for FMOD (rabbit IgG), 1:200 for PRG4 (mouse IgG), 1:250 for TNC (mouse IgG, sc-59884, Santa Cruz Biotechnology, Dallas, Texas), and 1:100 for ELN (mouse IgG, ab9519, Abcam, Cambridge, UK). Antibodies for DCN and PRG4 were a kind gift from Prof. Caterson, Cardiff University, UK, and the FMOD antibody was kindly provided by Prof. Roughley, McGill University, Canada. The secondary antibody incubation was performed with the Zytochem Plus HRP Polymer anti-rabbit for FMOD and anti-mouse for DCN, PRG4, TNC, and ELN (ZUC032-006 and ZUC050-006, Zytomed Systems, Berlin, Germany). Immunohistochemical staining was graded from 0 to 3 (low to high) on blinded sections, assessing stained area and staining intensity for DCN, FMOD, TNC, and ELN. For PRG4, where staining was confined to the pericellular area, staining intensity was measured using TissueFaxs (Tissuegnostics).

### Quantification of tendon elastin

The elastin content of SDFT and CDET samples from each age group (n=12; 3 from each age group) was quantified using the FASTIN™Elastin Assay (Biocolor, Carrickfergus, UK)(Godinho et al., 2017). Briefly, SDFT and CDET tissue was powdered (∼15 mg wet weight) and incubated with 750 µl of 0.25 M oxalic acid at 100 °C for 2 one hour cycles to extract all soluble a-elastin from the tissue. Preliminary tests showed two extractions were sufficient to solubilise all a-elastin from developing SDFT and CDET. Following extraction, samples and standards were processed in duplicate according to the manufacturer’s instructions and their absorbance determined spectrophotometrically at 513 nm (Spectrostar Nano microplate reader, BMG Labtech, Aylesbury, UK). A standard curve was used to calculate the samples’ elastin concentration and elastin was expressed as a percentage of tendon wet weight.

### Laser-capture microdissection

Laser-capture microdissection was used to collect samples from the fibre and matrix phase of SDFT samples from all age groups (n=4 for each age group with the exception of the 0-1 month group where n=3). For this purpose, 12 µm transverse cryosections were cut from the SDFT samples and mounted on steel frame membrane slides (1.4 µm PET membrane, Leica Microsystems, Wetzlar, Germany). Frozen sections were dehydrated in 70% and 100% ice-cold ethanol, allowed to briefly dry, and regions of fibre and matrix phase laser-captured on an LMD7000 laser microdissection microscope (Leica Microsystems, Wetzlar, Germany) and collected in LC/MS grade water (FisherScientific, Hampton, New Hampshire). Collected samples were immediately snap frozen and stored at −80°C for mass spectrometry analysis.

### Mass spectrometry analysis

Mass spectrometry analysis of laser-captured SDFT matrix and fibre phase samples was carried out as previously described by Thorpe et al. (2016)(Chavaunne T. Thorpe, Peffers, et al., 2016). Samples were digested for mass spectrometry analysis with incubation in 0.1% (w/v) Rapigest (Waters, Herts, UK) for 30 min at room temperature followed by 60 min at 60 °C and subsequent trypsin digestion. LC MS/MS was carried out at the University of Liverpool Centre for Proteome Research using an Ultimate 3000 Nano system (Dionex/Thermo Fisher Scientific, Waltham, Massachusetts) for peptide separation coupled online to a Q-Exactive Quadrupole-Orbitrap mass spectrometer (Thermo Scientific, Waltham, Massachusetts) for MS/MS acquisition. Initial ranging runs on short gradients were carried out to determine the sample volume to be loaded on the column and subsequently between 1-9 µL of sample was loaded onto the column on a one hour gradient with an inter-sample 30 min blank.

### Protein identification and label-free quantification

Fibre and matrix phase proteins were identified using Peaks® 8.5 PTM software (Bioinformatics Solutions, Waterloo, Canada), searching against the UniHorse database (http://www.uniprot.org/proteomes/). Search parameters used were: peptide mass tolerance 10 ppm, fragment mass tolerance 0.01 Da, fixed modification carbamidomethylation, variable modifications methionine oxidation and hydroxylation. Search results for peptide identification were filtered with a false discovery rate (FDR) of 1%, and for protein identification with a minimum of 2 unique peptides per protein, and a confidence score >20 (- 10lgp>20). Label-free quantification was also carried out using Peaks® 8.5 PTM software for the SDFT fibre and matrix phase separately. Protein abundances were normalised for collected laser-capture area and volume loaded onto the mass spectrometry column and differentially abundant proteins between the age groups in the SDFT fibre and matrix phase were identified using a fold change ≥2 and *p*<0.05 (PEAKS adjusted p values). The mass spectrometry proteomics data have been deposited to the ProteomeXchange Consortium via the PRIDE partner repository, with the dataset identifier PXD012169 and 10.6019/PXD012169.

### Gene ontology and network analysis

The dataset of identified proteins in the SDFT fibre and matrix phase were classified for cell compartment association with the Ingenuity Pathway Analysis software (IPA, Qiagen, Hilden, Germany) and for matrisome categories with The Matrisome Project database(Hynes & Naba, 2012). Protein pathway analysis for the differentially abundant proteins between age groups in the SDFT fibre and matrix phase was carried out in IPA. Protein interactions maps were created in IPA allowing for experimental evidence and highly predicted functional links.

### Neopeptide identification

For neopeptide identification, mass spectrometry data was analysed using Mascot server (Matrix Science) with the search parameters: enzyme semiTrypsin, peptide mass tolerance 10 ppm, fragment mass tolerance 0.01 Da, charge 2+ and 3+ ions, and missed cleavages 1. The included modifications were: fixed carbamidomethyl cysteine, variables oxidation of methionine, proline, and lysine, and the instrument type selected was electrospray ionization-TRAP (ESI-TRAP). The Mascot-derived ion score was used to determine true matches (*p*<0.05), where *p* was the probability that an observed match was a random event. The peptide list was exported and processed with the Neopeptide Analyser, a software tool for the discovery of neopeptides in proteomic data(Peffers et al., 2017). Obtained neopeptide abundances for each sample were normalised for total peptide abundance for that protein and sample, and normalised neopeptide abundances were subsequently summed for each protein and the total neopeptide abundance analysed for differential abundance across the age groups in the SDFT fibre and matrix phase using *p*<0.05 and FDR 5% (ANOVA and Benjamini-Hochberg FDR).

### Relative mRNA expression

RNA extraction from whole SDFT and CDET was carried out followed by reverse transcription. Quantitative real-time PCR (qRT-PCR) was performed on an ABI7300 system (Thermo Fisher Scientific Waltham, Massachusetts) using the Takyon ROX SYBR 2X MasterMix (Eurogentec, Liege, Belgium). qRT-PCR was undertaken using gene-specific primers for DCN, FMOD, BGN, COMP, COL1A1, COL1A2, COL3A1, TGFB1, and GAPDH as a reference gene previously validated (Supplementary Table 2). Relative expression levels were normalised to GAPDH expression and calculated with the formula E_−ΔCt_ following primer efficiency calculation.

### Statistical analysis

Statistical analysis was carried out in SigmaPlot (Systat Software Inc, San Jose, California) unless otherwise stated. Details of the n numbers for each experiment and the statistical test used for the analysis of the data are listed in Supplementary Table 3. Heatmaps were designed in GProX(Rigbolt et al., 2011). The Central Limit Theorem (CLT) was used to assume normality where n>30 and where n<30 normality was tested using the Shapiro-Wilks test. If data were found not to be normally distributed their log10 transformation or ANOVA on Ranks was used for statistical analysis but the original data was presented in graphs.

## Acknowledgements

This work was funded by the Horserace Betting Levy Board, PRJ/776. We would like to acknowledge the equine practices that provided samples for this study.

## Author Contributions

DEZ designed and performed experiments, analysed the data, and wrote the manuscript. CTT assisted with study design, data analysis, and edited the manuscript. YAK assisted with data collection and analysis. HLB, HRCS, PDC conceived the study, assisted with data analysis, and edited the manuscript.

## Data and materials availability

The mass spectrometry proteomics data have been deposited to the ProteomeXchange Consortium via the PRIDE partner repository, with the dataset identifier PXD012169 and 10.6019/PXD012169.

## Supplementary data

**Supplementary Figure 1.**
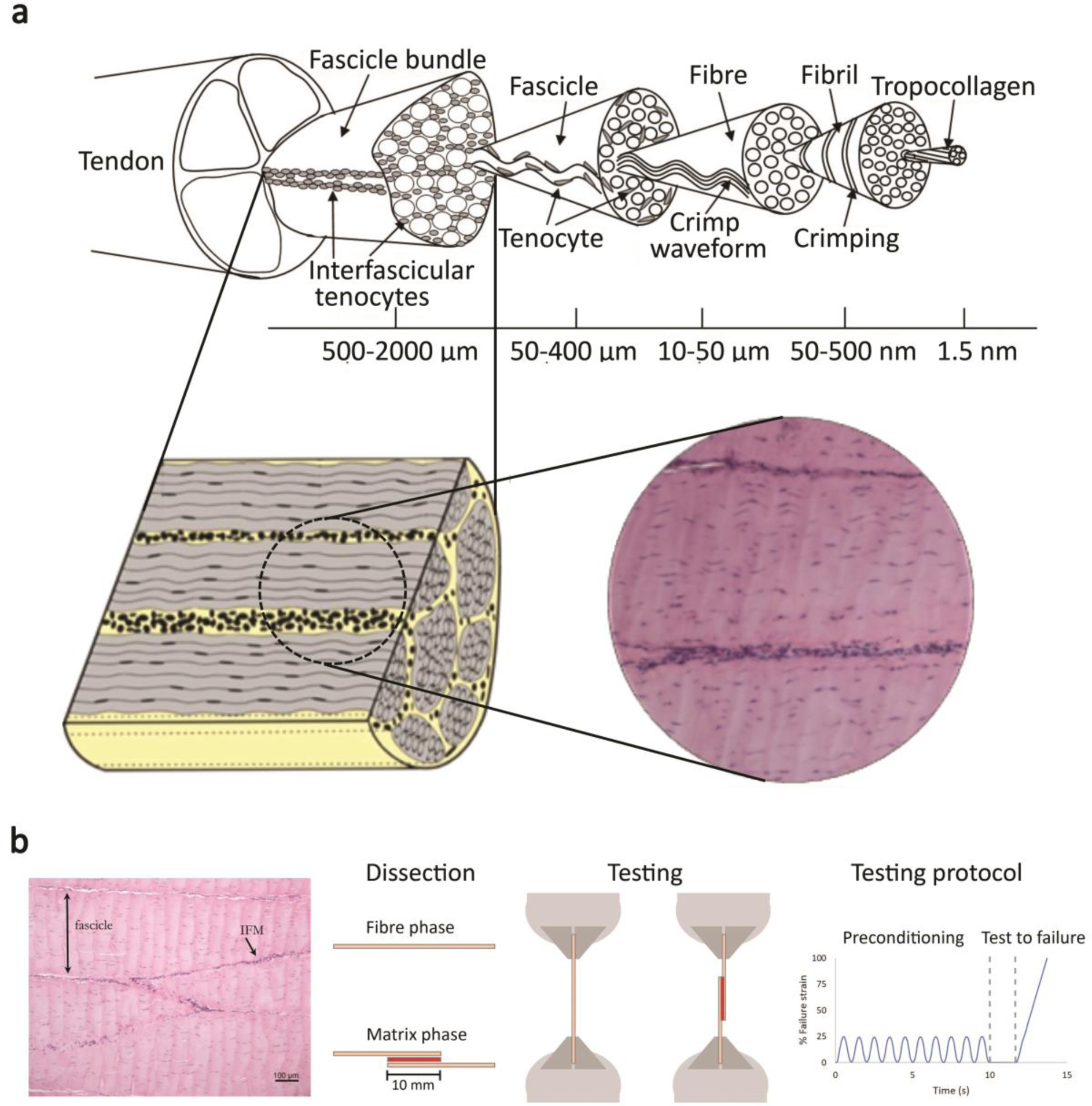
Tendon structure and graphical abstract for biomechanical testing. (**a**) Tendon structure (adapted from Spiesz et al. 2015)(Spiesz et al., 2015). (**b**) H&E section of fibre and matrix phase and schematic of fibre and matrix phase dissection and biomechanical testing.

**Supplementary Figure 2.**
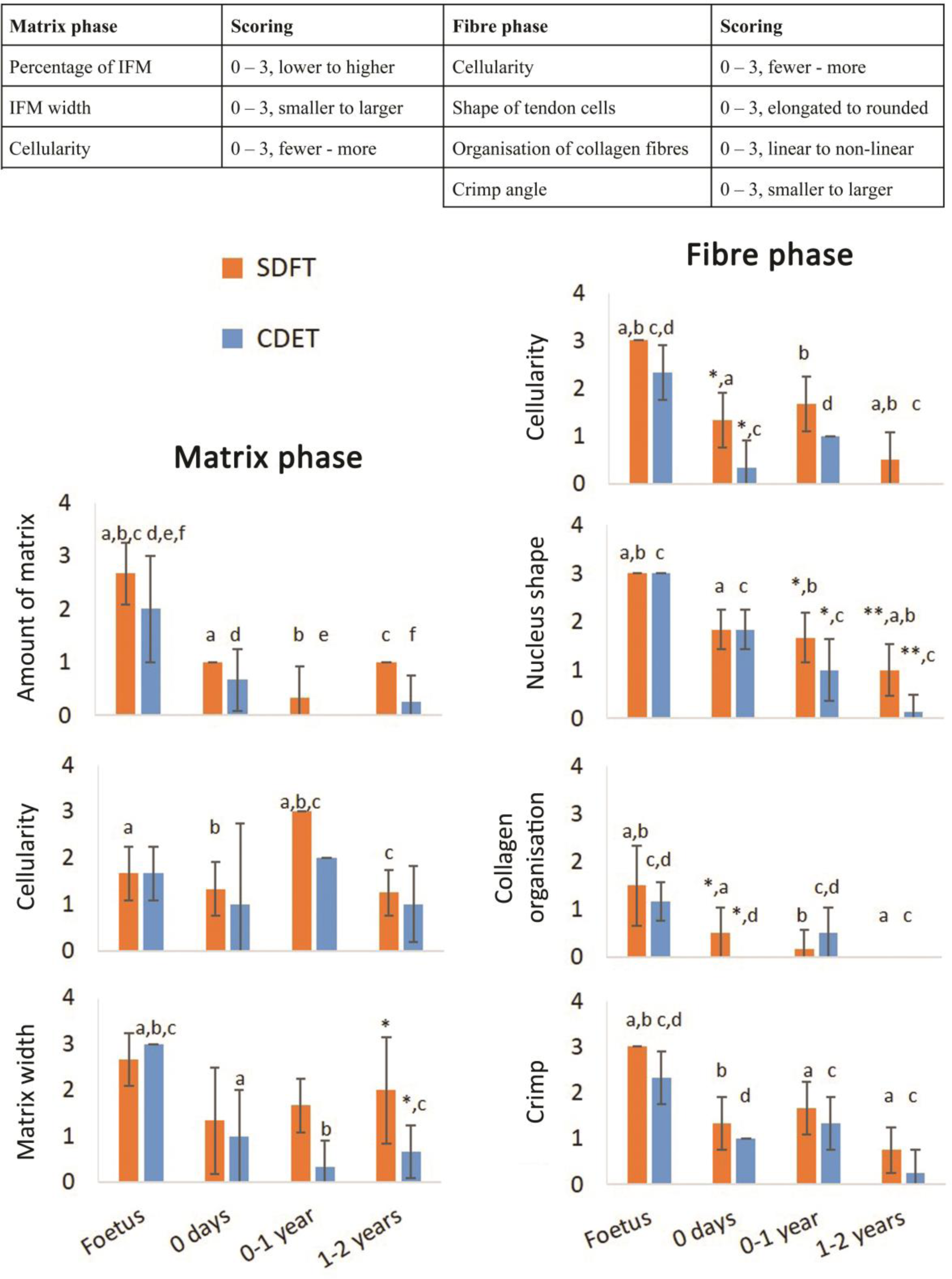
Scoring of histologic variables for the matrix and fibre phases in the SDFT and CDET through postnatal development. * significant difference between tendons, a-f significant difference between age groups. Error bars depict standard deviation.

**Supplementary Figure 3.**
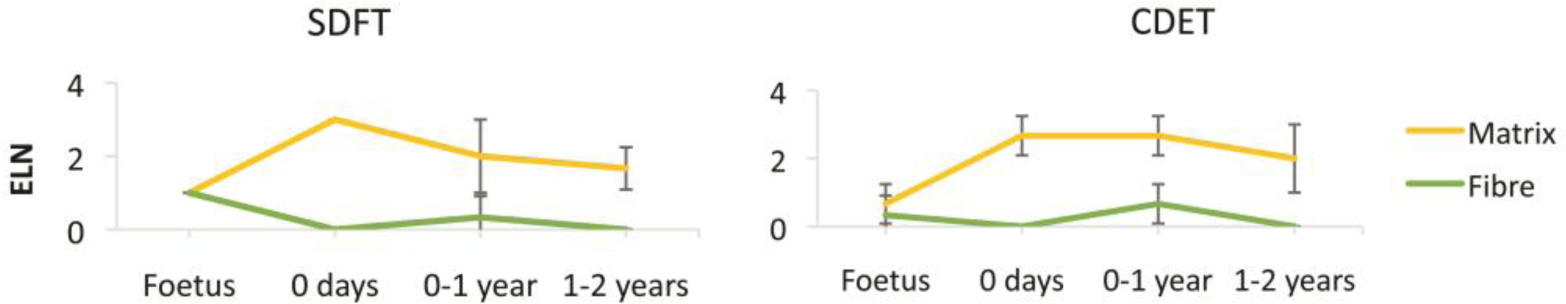
Scoring of ELN staining for the matrix and fibre phases in the SDFT and CDET through postnatal development. Error bars depict standard deviation.

**Supplementary Figure 4.**
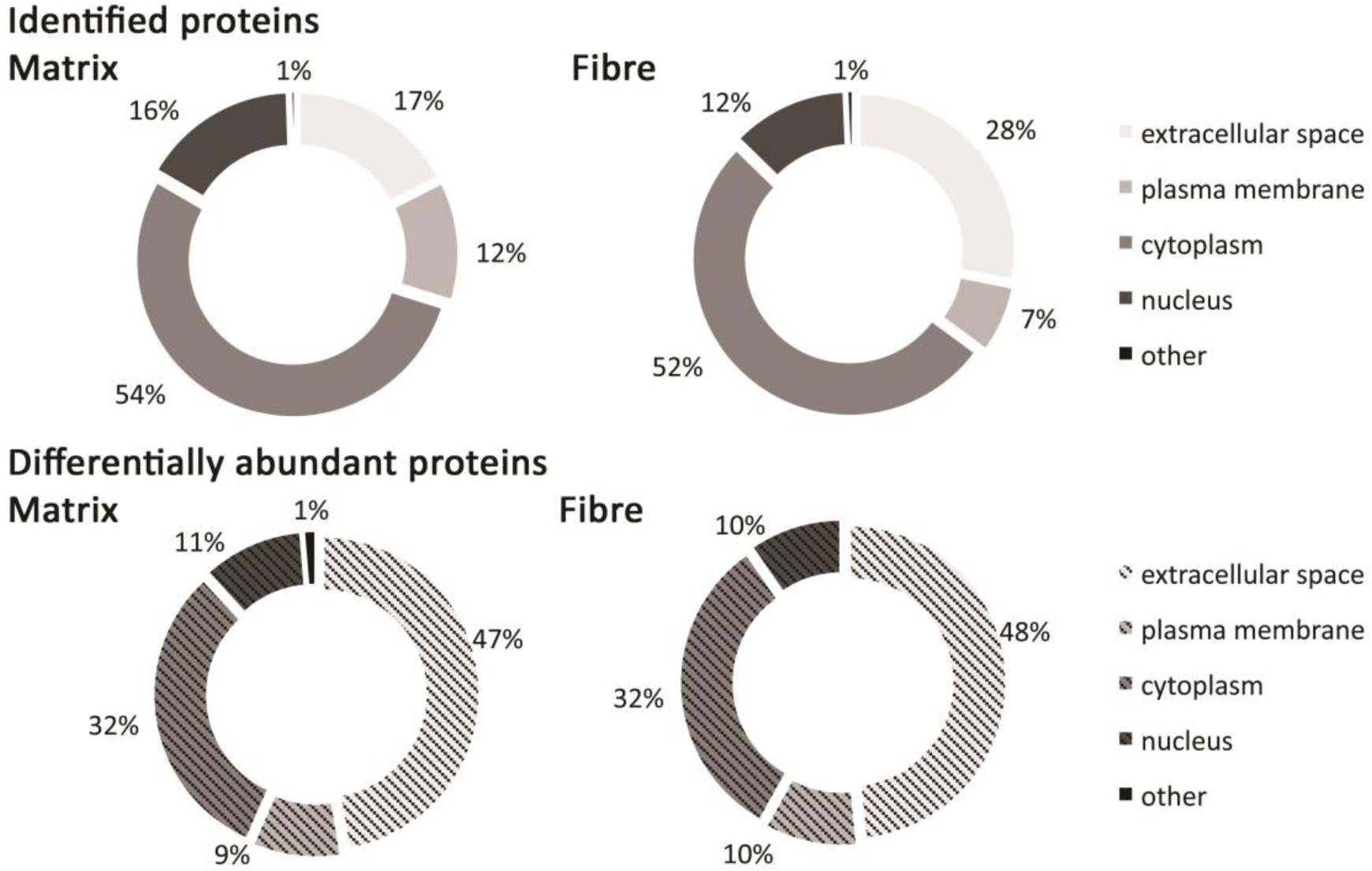
Classification of SDFT matrix and fibre phase identified proteins and differentially abundant proteins (*p*<0.05, fold change≥2) according to their associated location.

**Supplementary Figure 5.**
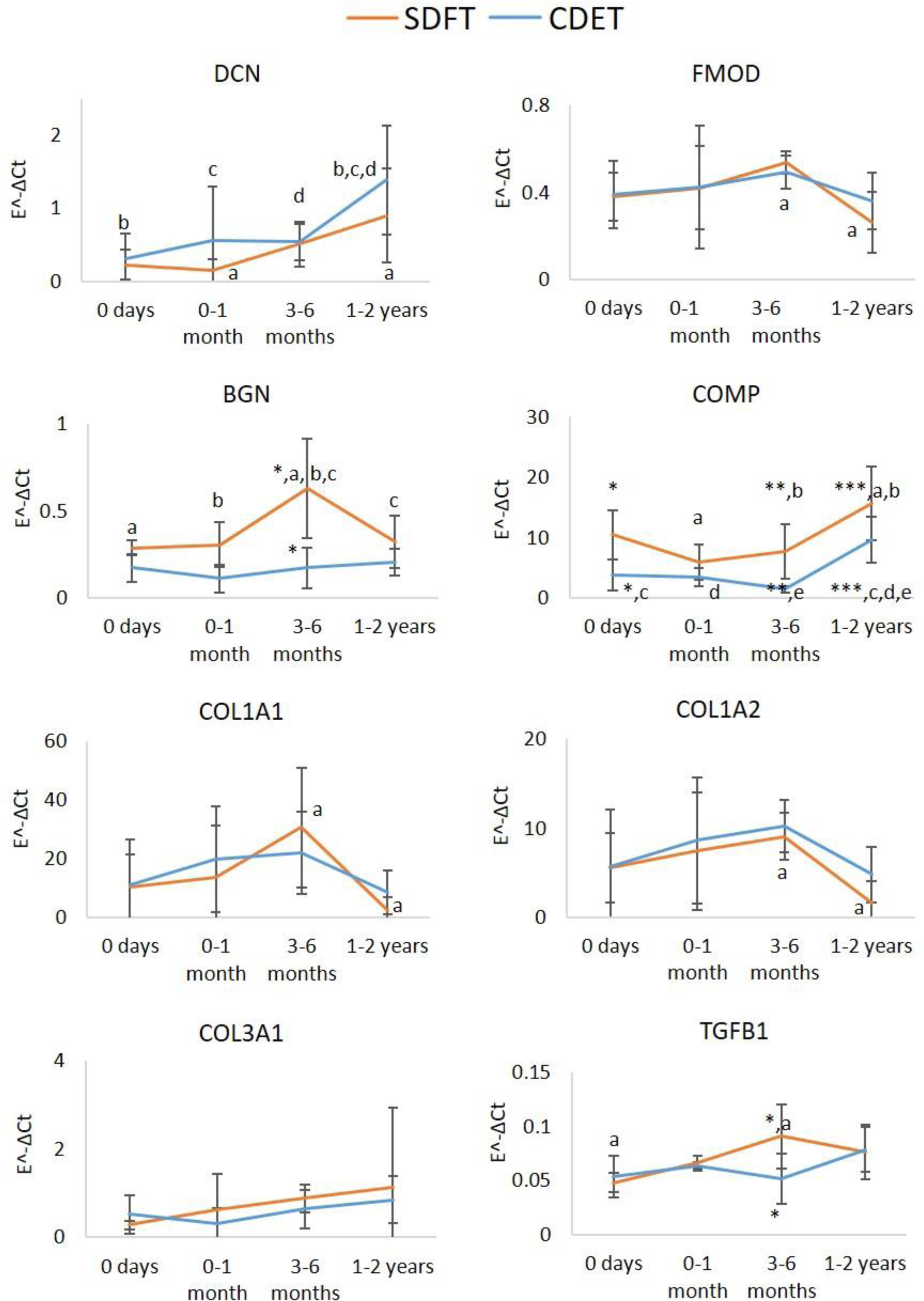
Relative mRNA expression of major ECM genes in whole tissue SDFT and CDET through postnatal development. * significant difference between tendons, a-e significant difference between age groups. Error bars depict standard deviation.

**Supplementary Table 1.**
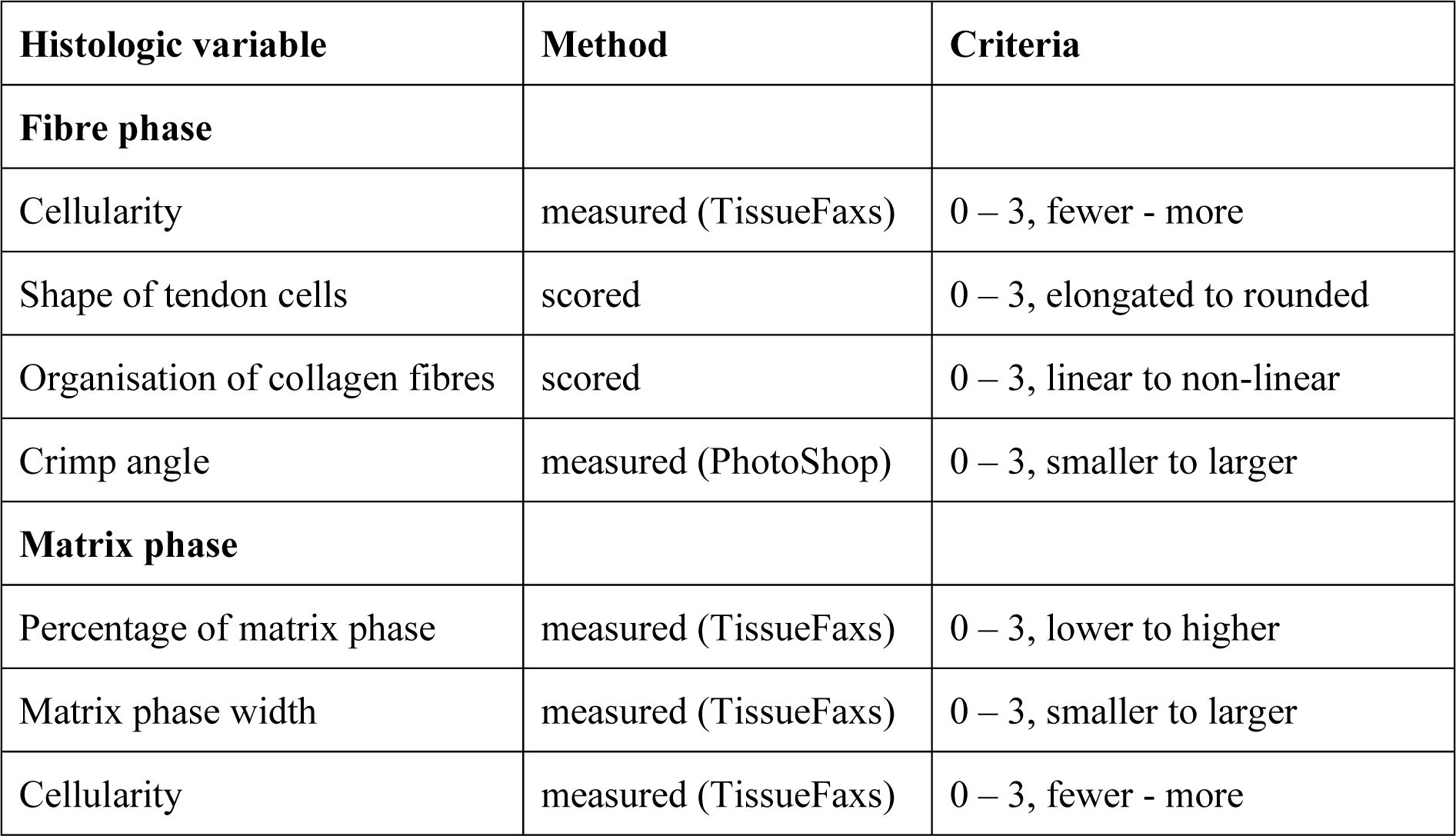
Histologic variables used in the H&E scoring of the SDFT and CDET sections and the analysis method and reporting criteria adopted. When software was adopted to acquire a parameter, the software is additionally reported. “Scored” is used to denote parameters analysed by blinded investigators.

**Supplementary Table 2.**
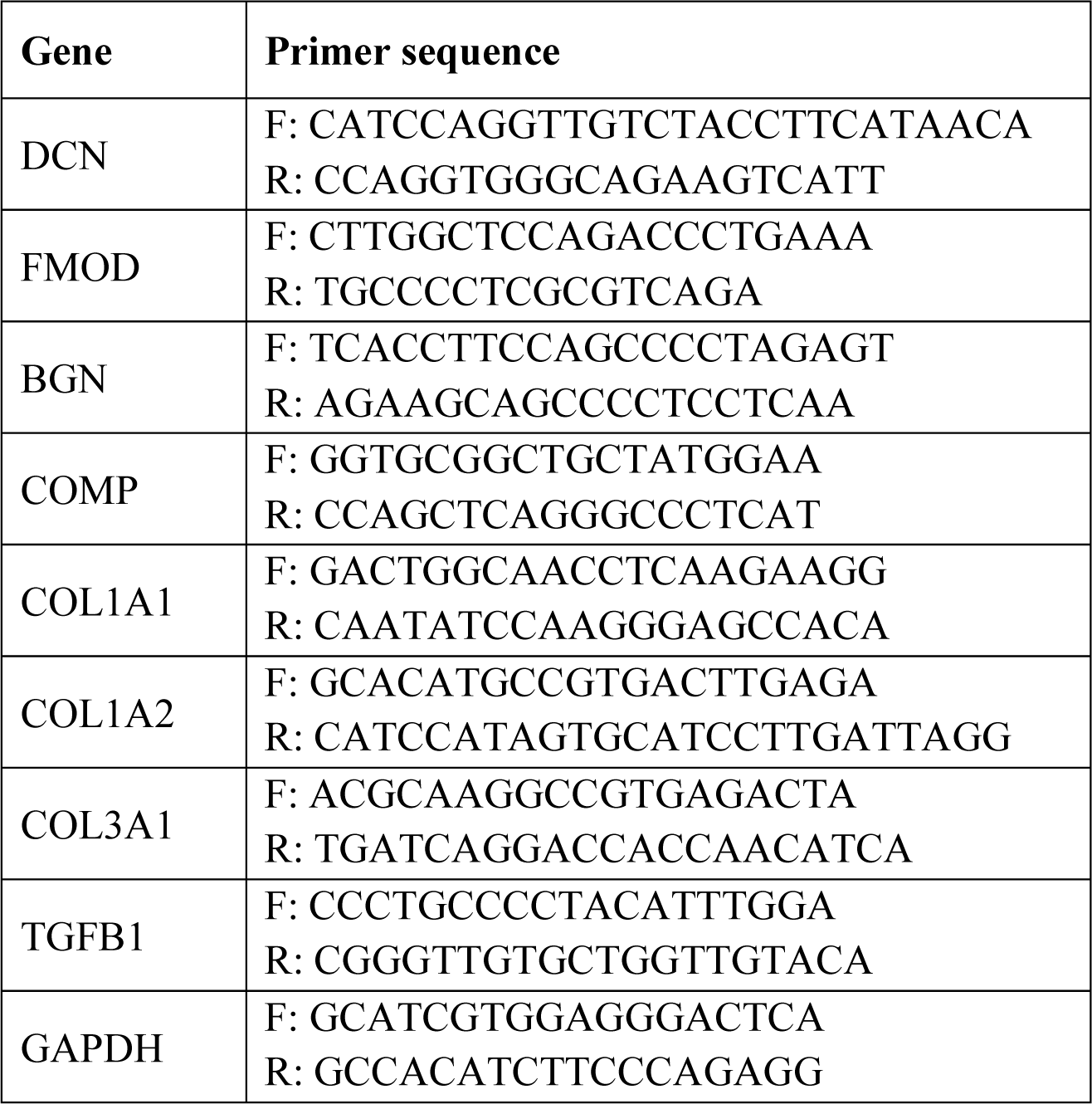
Gene primer sequences used in relative mRNA expression analysis.

**Supplementary Table 3.** Samples used for analysis along with statistical test used for analysis. n numbers represent biological replicates.

| Analysis | Tendon | Foetus | 0 days | 0-1 month | 3-6 months | 1-2 years | Statistical analysis |
| --- | --- | --- | --- | --- | --- | --- | --- |
| **Biomechanics** | SDFT fibre | n=4 | n=4 | n=3 | n=4 | n=4 | Two-way ANOVA. Outliers < or > mean ± 2xSD were removed. The average of 4-7 technical replicates was used (depending on sample availability). |
|  | SDFT matrix | n=4 | n=4 | n=3 | n=4 | n=4 |  |
|  | CDET fibre | n=4 | n=4 | n=3 | n=4 | n=4 |  |
|  | SDFT matrix | n=3 | n=4 | n=3 | n=4 | n=4 |  |
| **Histology and Immunohistochemistry** | SDFT | n=3 | n=3 | n=3 (0-1 year) |  | n=4 (hist.), 3 (IHC) | Kruskal-Wallis, Rank Sum Test |
|  | CDET | n=3 | n=3 | n=3 (0-1 year) |  | n=4 (hist.), 3 (IHC) |  |
| **Elastin quantification** | SDFT | n=3 | n=3 | n=3 (0-1 year) |  | n=3 | Two-way ANOVA. The average of 2 technical replicates was used. |
|  | CDET | n=3 | n=3 | n=3 (0-1 year) |  | n=3 |  |
| **Proteomics** | SDFT fibre | n=3 | n=4 | n=3 | n=4 | n=4 | Differentially abundant proteins – PEAKS Q |
|  | SDFT matrix | n=4 | n=4 | n=3 | n=4 | n=4 | Differential total neopeptide abundance – One-way ANOVA, Benjamini-Hochberg correction |
| **Relative mRNA expression** | SDFT | - | n=4 | n=3 | n=4 | n=4 | Two-way ANOVA |
|  | CDET | - | n=4 | n=3 | n=4 | n=4 |  |

**Supplementary Table 4.** Collagens and proteoglycans identified in SDFT matrix and fibre phases. (1% FDR, protein −10lgP >20, and ≥2 unique peptides).

| Collagens | Matrix | Fibre | Proteoglycans | Matrix | Fibre |
| --- | --- | --- | --- | --- | --- |
| COL1A1 | + | + | ACAN | + | + |
| COL1A2 | + | + | ASPN | + | + |
| COL2A1 | + | + | BGN | + | + |
| COL3A1 | + | + | CHAD | - | + |
| COL4A1 | + | - | DCN | + | + |
| COL4A2 | + | - | FMOD | + | + |
| COL4A3 | - | + | HAPLN1 | + | + |
| COL4A6 | + | - | HSPG2 | + | + |
| COL5A1 | + | + | KERA | + | + |
| COL5A2 | + | + | LUM | + | + |
| COL5A3 | + | - | OGN | + | + |
| COL6A1 | + | + | PRELP | + | + |
| COL6A2 | + | + | VCAN | + | - |
| COL6A3 | + | + |  |  |  |
| COL8A1 | + | - |  |  |  |
| COL11A1 | + | + |  |  |  |
| COL12A1 | + | + |  |  |  |
| COL14A1 | + | + |  |  |  |
| COL15A1 | + | + |  |  |  |
| COL17A1 | - | + |  |  |  |
| COL18A1 | + |  |  |  |  |
| COL21A1 | + | + |  |  |  |
| COL28A1 | + |  |  |  |  |

